# CCL2/CCR2 inhibition in atherosclerosis: a meta-analysis of preclinical studies

**DOI:** 10.1101/2021.04.16.439554

**Authors:** Luka Živković, Yaw Asare, Jürgen Bernhagen, Martin Dichgans, Marios K. Georgakis

## Abstract

**Rationale:** The CC-chemokine ligand-2 (CCL2)/ CC-chemokine receptor-2 (CCR2) axis governs monocyte recruitment to atherosclerotic lesions. Coherent evidence from experimental studies employing genetic deletion of CCL2 or CCR2 and human epidemiological studies support a causal involvement of the CCL2/CCR2 axis in atherosclerosis. Still, preclinical studies testing pharmacological inhibition of CCL2 or CCR2 in atheroprone mice apply widely different approaches and report inconsistent results, thus halting clinical translation.

**Objective:** To systematically review and meta-analyze preclinical studies pharmacologically targeting the CCL2/CCR2 axis in atherosclerosis in an effort to inform the design of future trials.

**Methods and Results:** We identified 14 studies testing CCL2/CCR2 inhibition using 11 different pharmacological agents in mouse models of atherosclerosis. In meta-analyses, blockade of CCL2 or CCR2 attenuated atherosclerotic lesion size in the aortic root or arch (*g*=-0.75 [-1.17 to -0.32], p=6×10^−4^; N=171/171 mice in experimental/control group), the carotid (*g*=-2.39 [-4.23 to -0.55], p=0.01; N=24/25) and the femoral artery (*g*=-2.38 [-3.50 to -1.26], p=3×10^−5^; N=10/10). Furthermore, CCL2/CCR2 inhibition reduced intralesional macrophage accumulation and increased smooth muscle cell content and collagen deposition, consistent with a plaque-stabilizing effect. While there was heterogeneity across studies, the effects of CCL2/CCR2 inhibition on lesion size correlated with reductions in plaque macrophage accumulation, in accord with a prominent role of CCL2/CCR2 signaling in monocyte recruitment. Subgroup analyses revealed similar efficacy of both CCL2- and CCR2-inhibiting approaches across different atherosclerosis models in reducing lesion size and intralesional macrophage accumulation, but stronger atheroprotective effects in carotid and femoral arteries, as compared to the aorta.

**Conclusions:** Pharmacological targeting of CCL2 or CCR2 lowers atherosclerotic lesion burden and confers plaque stability in mice across different vascular territories, drug candidates, and models of atherosclerosis. Our findings in conjunction with recent human data highlight the translational potential of targeting the CCL2/CCR2 axis in atherosclerosis and can inform future clinical trials.

**Subject codes:** atherosclerosis, inflammation, vascular biology, translational studies

## Introduction

Cardiovascular disease remains the leading cause of mortality and disability.^1, 2^ Multiple lines of experimental and clinical evidence implicate inflammatory mechanisms in atherosclerosis, the predominant pathology underlying cardiovascular disease. Recent clinical trials have provided proof-of-concept for the role of inflammation in atherosclerosis by demonstrating the potential of anti-inflammatory therapies to lower cardiovascular risk.^3-5^ Specifically, canakinumab^3^ and colchicine^4, 6^ were found to lower the risk of recurrent cardiovascular events in patients with a history of coronary artery disease. While interventional studies in humans have so far mostly focused on the inflammasome-interleukin-1β (IL-1β)-interleukin-6 (IL-6) axis,^7^ recent experimental and epidemiological studies place emphasis on other mediators of inflammation, as has specifically been shown for the chemokine system.^8-10^ Targeting alternative inflammatory pathways with a more specific role in atherosclerosis could increase efficacy and improve the safety profile of anti-inflammatory approaches, thus moving them closer to clinical translation.

CC-motif chemokine ligand 2 (CCL2), is one of the first CC family chemokine described and implicated in atherosclerosis.^11-13^ CCL2 primarily acts by binding to CC-chemokine receptor 2 (CCR2) on the surface of classical monocytes, thus mobilizing them from the bone marrow to the circulation and attracting them to sites of inflammation^14^ including the arterial subendothelium.^11, 15-17^ CCL2/CCR2 signaling governs rolling and adhesion of monocytes on the endothelial lining of atherosclerotic lesions.^18^ Hyperlipidemic atheroprone mice deficient for either *Ccl2*^13^ or *Ccr2*^12, 19^ exhibit substantial reductions in the number and size of atherosclerotic lesions, as well as reductions in lipid deposition and macrophage accumulation in the arterial walls, thus supporting a causal role of the CCL2/CCR2 pathway in atherogenesis.

The potential importance of these findings for the development of therapeutic strategies is illustrated by recent studies demonstrating a causal role of CCL2 in human atherosclerosis. First, using a Mendelian randomization approach, we recently found higher genetically proxied circulating levels of CCL2 to be associated with a higher risk of ischemic stroke, in particular large artery stroke, and a higher risk of coronary artery disease and myocardial infarction.^20^ Second, higher measured levels of circulating CCL2 were associated with a higher risk of ischemic stroke, coronary artery disease, and cardiovascular mortality in population-based cohorts of individuals free of cardiovascular disease at baseline.^21, 22^ Third, CCL2 levels quantified in atherosclerotic plaques from individuals undergoing carotid endarterectomy showed significant associations with histopathological, clinical, and molecular features of plaque vulnerability.^23^

While these studies identify the CCL2/CCR2 axis as a promising pharmacological target for the treatment of atherosclerosis, there are only limited data from randomized trials specifically targeting this pathway in the context of human atherosclerosis. In a study on 108 patients with cardiovascular risk factors and high circulating levels of high-sensitivity C-reactive protein (hsCRP), those treated with a single intravenous infusion of MLN1202, a humanized monoclonal antibody against CCR2, exhibited significant reductions in hsCRP levels after 4 weeks and continuing through 12 weeks after dosing.^24^ However, this phase II trial was not designed to investigate clinical endpoints.

Preclinical studies have explored various pharmacological approaches targeting the CCL2/CCR2 axis in models of atherosclerosis. Still, these studies reported largely variable and partly inconsistent results, possibly reflecting differences in the properties of the individual pharmacological agents, the selected drug targets (CCL2 or CCR2), the molecular sites in their structures targeted by the agents, the animal models under study, lesion stages at initiation of the intervention, duration of treatment, and the vascular beds under examination. Against this background, we aimed to analyze the available evidence from preclinical studies testing pharmacological inhibition of the CCL2/CCR2 pathway in atherosclerosis-prone mice and quantify the effects of the interventions on lesion size and features of plaque stability (macrophage accumulation, smooth muscle cell content, and collagen deposition). We further aimed to detect potential sources of heterogeneity of their efficacy including those related to pharmacological properties of the inhibitor, vascular bed, animal model, duration of treatment, and stage of atherosclerosis. We therefore performed a systematic review and meta-analysis of preclinical studies in an effort to inform the design of future clinical trials in humans.

## Methods

### Search strategy

The SyRF 9-step outline and SYRCLE’s protocol template for conception of animal meta-analyses^25^ were used to design the search strategy, study and outcome selection, and statistical processing of the extracted data for this systematic review and meta-analysis. To identify eligible articles, we screened PubMed from its inception to April 7^th^, 2021 without restrictions in language or publication year, using the following predefined search strategy: (“CC chemokine ligand 2” OR “C-C chemokine ligand 2” OR “C-C motif chemokine ligand 2” OR “C-C motif chemokine receptor type 2” OR “C-C chemokine receptor type 2” OR “CC chemokine receptor type 2” OR “monocyte chemoattractant protein” OR “monocyte chemotactic protein” OR CCL2 OR MCP-1 OR MCP1 OR CCR2) AND (atherogenesis OR atherosclerosis OR atheroprogression OR atherosclerotic OR plaque OR stroke OR ((cardiovascular OR ischemic OR cerebrovascular OR coronary) AND disease) OR (myocardial AND infarction)). A published search filter was employed to limit displayed entries to those referring to animal experiments.^26^ Additionally, the reference lists of all eligible studies were screened. Eligible articles were evaluated for potential overlap of data. One reviewer (L.Z.) performed the initial screening and all potentially eligible articles were further independently screened by an additional reviewer (M.G.); differences were resolved in consensus.

### Eligibility criteria

#### Population

Articles were deemed eligible if they described experimental inhibition of CCL2 or CCR2 in an *in vivo* mouse model of atherosclerosis. Specifically, eligible studies were required to use atherosclerosis-prone mouse models, such as *Apoe*^*-/-*^, *Ldlr*^*-/-*^, or ApoE3Leiden mice that were fed a normal chow or high-fat “Western-type” diet (WTD). Models of accelerated atherosclerosis following arterial injury in atherosclerosis-prone mice were also considered eligible.^27-29^ Studies referring to other animals beyond mice were not included in this review.

#### Intervention

Eligible studies had to explore the effects of a pharmacological intervention directly interfering with and inhibiting CCL2 or CCR2, such as orthosteric or allosteric receptor antagonists, competitive inhibitors of chemokine-receptor interaction, or antibodies. Studies that made use of gene therapy by means of transfecting plasmids, such as 7ND,^30-33^ were also considered eligible, provided that the encoded protein was a direct inhibitor of CCL2 or CCR2. Studies that examined pharmacological agents or nutritional compounds that indirectly downregulate the CCL2/CCR2 axis by interfering with upstream agents or downregulated CCL2 or CCR2 expression were deemed ineligible. Eligible studies also required a control group of animals which were injected with a vehicle or were fed an inhibitor-free diet.

#### Outcomes

Eligible studies needed to provide a quantified measurement of atherosclerotic plaque burden as an outcome. Histopathological quantification of lesion size/area, plaque size/area, neointimal area, or lipid-staining area following hematoxylin-eosin, Oil Red O, trichrome or pentachrome staining of vessel cross-sections were required for inclusion in the meta-analysis. Studies providing measurements of intima/media ratio were excluded, because these readouts fail to distinguish atherosclerotic plaque burden from intimal hyperplasia and vascular thickening. Apart from the well-established lesion quantification in the aortic root or arch, studies measuring carotid or femoral artery lesions were also included in the meta-analysis.

Additional predefined outcomes entailed plaque stability features including macrophage accumulation (expressed as Mac2/3-positive or Moma-2/3-positive content), smooth muscle cell content (expressed as smooth-muscle-actin-positive content), and collagen deposition. Plaque stability outcomes were only included if they were normalized to plaque size. We further explored the following as secondary outcomes: effects of the intervention on additional measurements that had not been predetermined, when explored by at least three individual studies. These included body weight, plasma cholesterol and triglyceride levels, circulating monocyte count, plasma CCL2 levels, and aortic expression of CCR2, IL-6, and TNF-α.

### Study quality assessment

We examined potential sources of bias with the SYRCLE risk of bias tool that was specifically designed for preclinical studies.^34^ The tool evaluates studies for selection bias (3 items), performance bias (2 items), detection bias (2 items), attrition bias (1 item), reporting bias (1 item), and other sources of bias (1 item). If the study did not provide any information whether a type of bias was appropriately addressed, the risk was classified as unclear, except for the detection bias item of outcome assessor blinding, where all studies that failed to report on blinding were ascribed a high risk of bias. Full texts, figure legends and supplementary materials were considered in the risk of bias assessment.

### Data abstraction

Absolute values, number of specimens, and either standard error or standard deviation in both intervention and control groups were extracted for each outcome. Where numerical data was not available, values were extracted from figures. Additionally, information pertaining to experimental setup such as inhibitor used and its target (CCL2 or CCR2), blood vessel under examination, mouse background and genetic model, sex, type of diet and (where applicable) start and duration of WTD feeding, start and duration of inhibitor administration, and additional pharmacological interventions were recorded for each experimental group. Where abstractable data were not presented in the published article or the supplementary materials, the corresponding author was contacted for providing the required information. One reviewer (L.Z.) performed the data abstraction and all data were further checked by a second reviewer (M.G.).

### Meta-analysis

For all studies and outcomes, we calculated standardized mean differences (Hedges’ *g*) between the experimental and the control groups using the Hedges’ approach to account for the small sample sizes.^35^ We then pooled the individual study estimates using DerSimonian-Laird random effects models to account for the expected heterogeneity between studies. For the main outcomes (lesion size and plaque stability characteristics), we performed the analysis separately for the different examined vessels (aortic root or arch, carotid artery, femoral artery). We calculated between-study heterogeneity with the *I*^*2*^ and the Cochran *Q* statistic. *I*^*2*^ exceeding 50% or 75% was considered as moderate and high heterogeneity, respectively.^36^ Finally, we performed Egger regression to explore potential small-study effects in our main analysis that would indicate presence of publication bias.^37^ Funnel plots were also created and visually inspected for asymmetry due to small-study effects.

To account for potential sources of heterogeneity, we performed a series of subgroup and meta-regression analyses. Specifically, we explored if the stage of lesion progression at the time of intervention start influenced the intervention effects. Lesion stage was classified as early, intermediate or advanced on the basis of mouse model, diet used, and age at intervention: *Apoe*^*-/-*^ and *Ldlr*^*-/-*^ mice fed a chow diet were assumed to exhibit early lesions until they were 15 weeks old, intermediate lesions between 15-20 weeks, and advanced lesions after 20 weeks of age. The resepective intervals in WTD-fed mice were until 10 weeks (early lesions); 10-15 weeks (intermediate lesions), and after 15 weeks (advanced lesion)s.^38-40^ We further performed subgroup analyses by target of intervention (CCL2 vs. CCR2), animal model (*Apoe*^-/-^ vs. *Ldlr*^-/-^), type of diet (chow vs. WTD) and sex. Finally, we carried out meta-regression analyses, exploring whether the effects of the interventions on plaque stability characteristics or other secondary outcomes, could explain heterogeneity in the effects on lesion size.

All analyses were performed using Stata 16.1 (College Station, United States).

## Results

The results of the search strategy are summarized in **Fig. 1**. The PubMed search returned 3,945 entries, out of which 279 articles were assessed for eligibility through inspection of their full texts. Of them, 16 articles met our eligibility criteria. Two of them presented data already available in another publication,^41, 42^ whereas one article^43^ did not present any abstractable data. One additional article^44^ was identified through screening the reference lists of the eligible studies. As such, a total of 14 articles^16, 30-33, 44-52^ were eventually deemed eligible for inclusion in our systematic review and meta-analysis.

**Figure 1.**
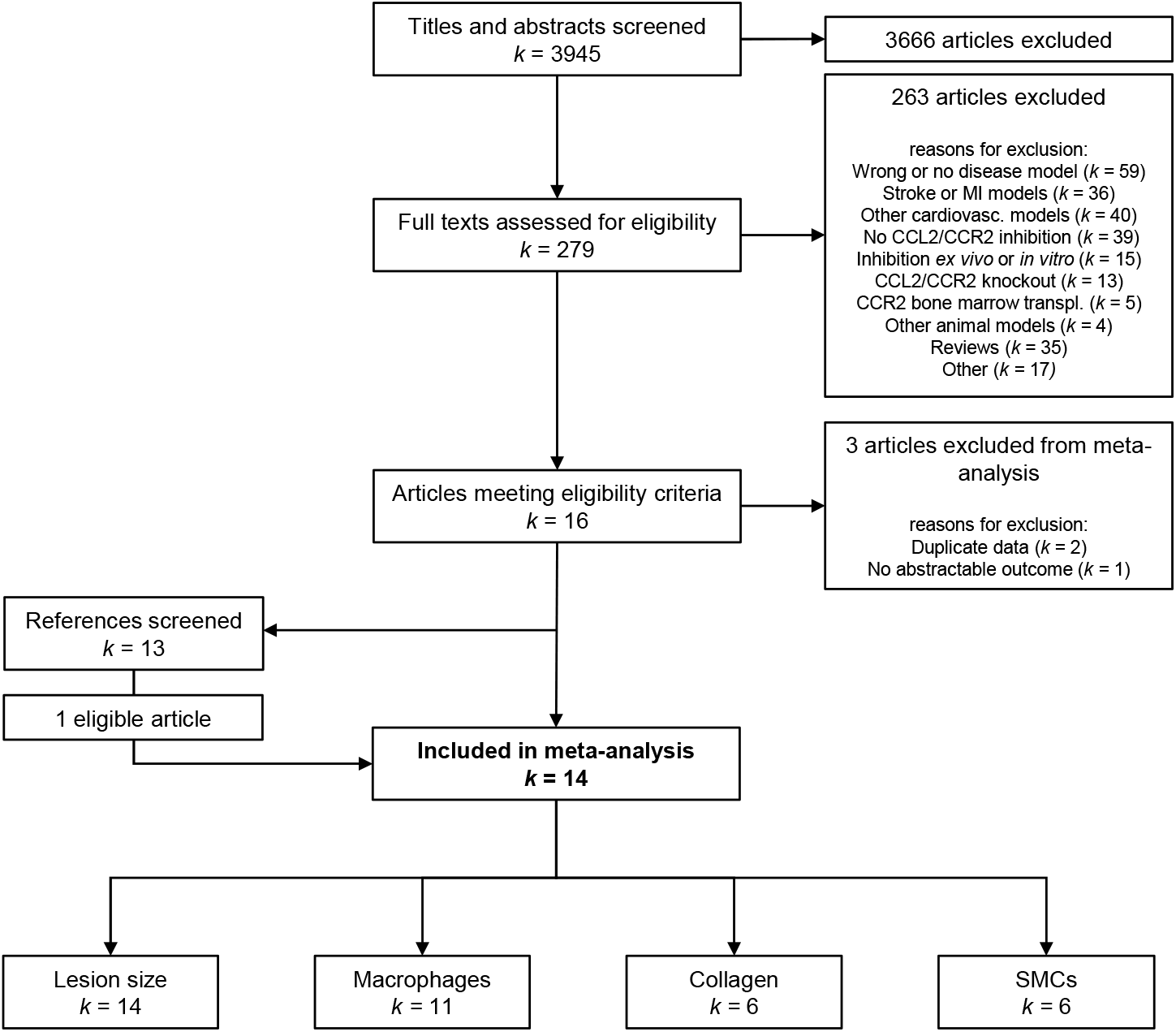
Flowchart of the study selection process. The search was performed in Medline through the PubMed engine and articles were evaluated for eligibility on the basis of their titles, abstracts, and full-texts. *k* = number of articles. SMCs = smooth-muscle cells.

### Characteristics of eligible studies

**Table 1** summarizes key characteristics of the included studies, which were published between 2001 and 2018. All eligible studies used hyperlipidemic mouse models of atherosclerosis. Specifically, 12 studies used *Apoe*^*–/–*^ mice, one *Ldlr*^*–/–*^ mice, and one ApoE3Leiden mice. Nine studies relied on WTD feeding in their experimental setup, whereas in four studies mice were fed normal chow, with one study not specifying any type of diet. Most of the studies that used a WTD-initiated high-fat feeding before the age of ten weeks. Timing of intervention and duration of treatment differed widely in the eligible studies ranging from four to 30 weeks (age at initiation of treatment) and three to 12 weeks (treatment duration), respectively.

**Table 1.**
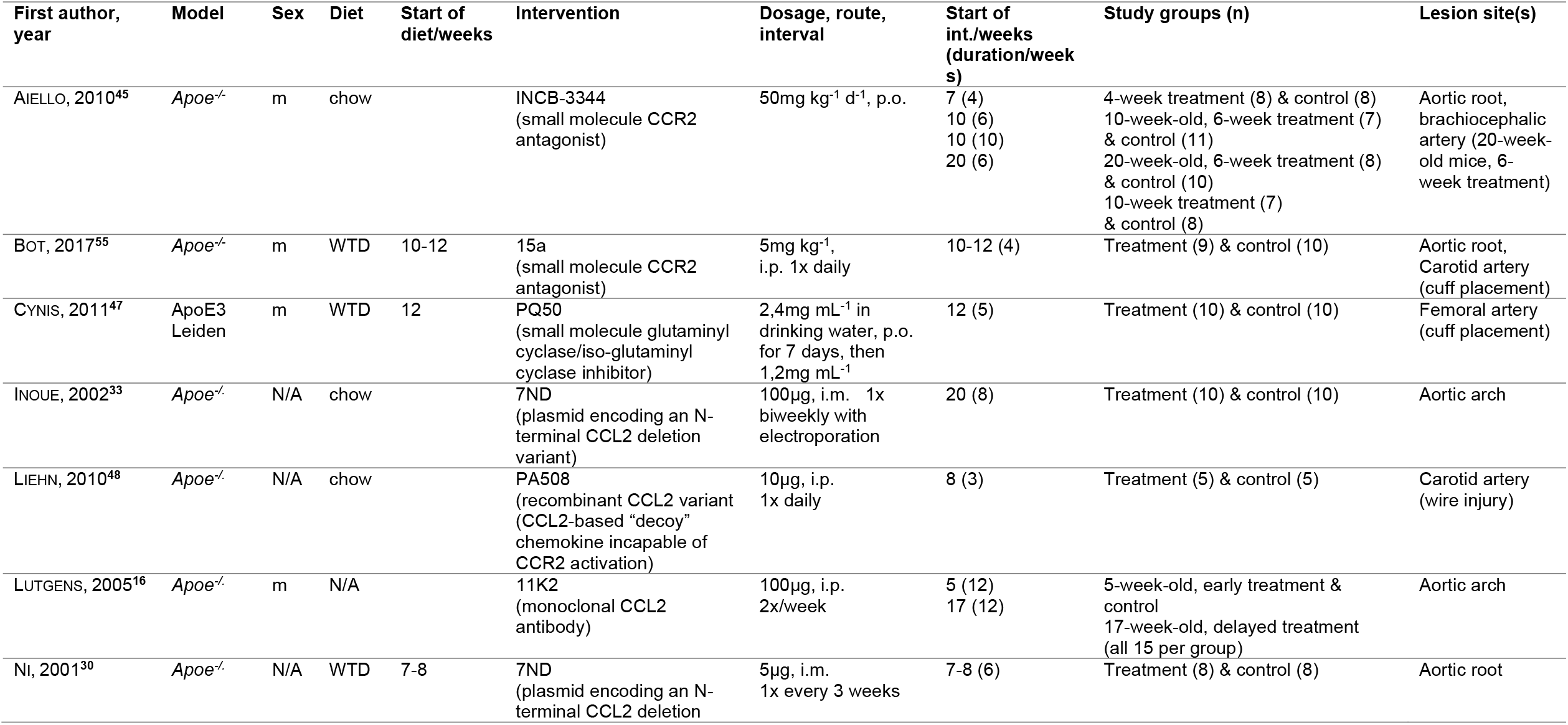
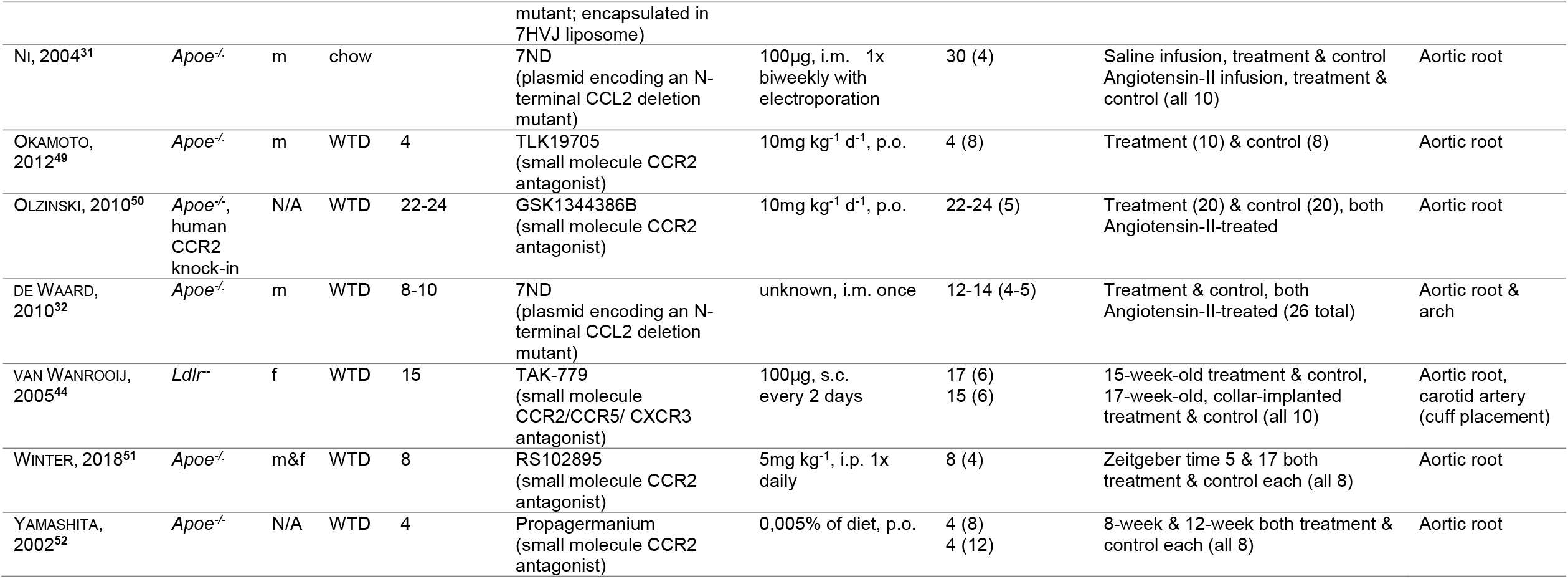
Descriptive characteristics of eligible studies included in the meta-analysis. N/A = Information not provided in the study; m = male, f = female; WTD = Western-type diet; d = day; p.o. = oral gavage, i.p. = peritoneal injection, i.m. = intramuscular injection, s.c. = subcutaneous injection.

Twelve studies targeted CCR2 with an inhibitory compound. Three studies used commercially available inhibitors.^44, 51, 52^ Specifically, Yamashita et al.^52^ tested Propagermanium, an organometallic, which has been used in the treatment of chronic hepatitis B,^53^ whereas Van Wanrooij et al. used TAK-779, a small molecule inhibitor of CCR2, CCR5, and CXCR3^46^ that is under examination as an HIV entry inhibitor.^44^ Winter et al.^51^ tested RS102895, a CCR2 small molecule antagonist^54^ in a chronopharmacological study. Four studies examined proprietary inhibitors that were developed in-house^49, 50, 55, 56^ and four studies utilized a plasmid encoding for an N-terminal deletion mutant of CCL2 (termed 7ND) as a therapeutic compound,^30-33^ which inhibits CCL2 signaling by functioning as a dominant-negative inhibitor of CCL2.^57, 58^ Finally, Liehn et al.^48^ opted for a similar approach using a recombinant N-terminal truncate of CCL2. Only two studies used approaches directly targeting CCL2: Lutgens et al.^16^ employed a monoclonal anti-CCL2 antibody, whereas Cynis et al.^47^ used a small molecule blocking essential posttranslational modifications on the N-terminus of CCL2. Regarding study outcomes, all but two studies examined plaque burden either in the aortic root or arch, whereas three studies explored lesions in the carotid artery. Cynis et al.^47^ opted for a femoral artery wire injury model in ApoE3Leiden mice, where lesion development was artificially induced and accelerated. Similarly, Liehn et al.^48^ conducted wire injury in the carotid artery of *Apoe*^*-/-*^ mice.

### Inhibiting CCL2 or CCR2 reduces lesion size and skews plaques towards a stable phenotype

Twenty-two treatment arms from all 14 studies were included in the meta-analysis for lesion size, as displayed in **Fig. 2**. Blockade of CCL2 or CCR2 resulted in a significant decrease in atherosclerotic lesion size in the aortic root or arch (*g*=-0.75 [-1.17 to -0.32], p=6×10^−4^), as derived after pooling 18 study arms (171 animals in experimental group, 171 controls). Significant decreases were also found in both the carotid (*g*=-2.39 [-4.23 to -0.55], p=0.01, k= 3 study arms, 24 animals in experimental group, 25 controls) and femoral arteries (*g*=-2.38 [-3.50 to -1.26], p=3×10^−5^, k= 1 study arm, 10 animals in experimental group, 10 controls). There was a significant difference in the effects of CCL2/CCR2 inhibition across the three vascular beds (p=0.01) with larger effects seen in the carotid and femoral arteries, as compared to the aortic root and arch.

**Figure 2.**
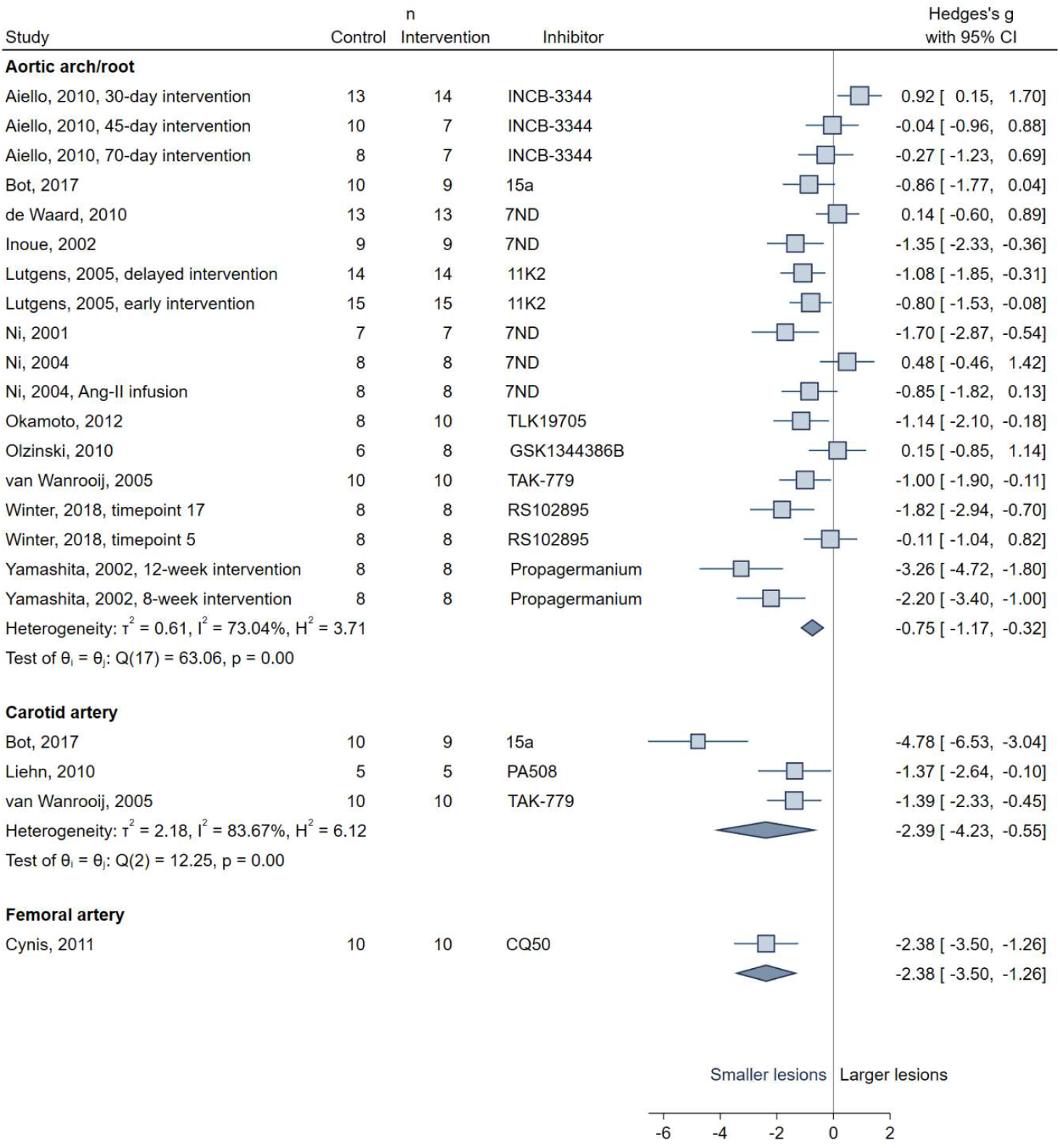
Forest plot of the effects of CCL2/CCR2 inhibition versus control (Hedges’ g) on atherosclerotic lesion size in the aortic arch and root, the carotid, and the femoral artery. Shown are the standardized mean differences, calculated as Hedges’ *g*, with their respective 95% confidence intervals per study. Plot squares are weighted for study size and correspond to individual effects, whereas plot whiskers correspond to the 95% confidence intervals. Diamonds indicate the pooled effects for each vascular bed. τ^2^, I^2^ and H^2^ as indicators of group heterogeneity as well as test of group homogeneity θi – θj are displayed for the groups containing more than one study.

CCL2/CCR2 inhibition further reduced the intralesional macrophage accumulation in the aortic arch and root (*g*=-0.76 [-1.11 to -0.41], p=2×10^−5^, k= 12 study arms, 112 animals in experimental group, 111 controls) (**Fig. 3**), while leading to an increase in collagen deposition (*g*=0.70 [0.16 to 1.24], p=0.011, k= 6 study arms, 60 animals in experimental group, 60 controls) and smooth-muscle cell content (*g*=0.95 [0.24 to 1.66], p=0.009, k= 6 study arms, 61 animals in experimental group, 61 controls), consistent with a more stable plaque phenotype.^59^

**Figure 3.**
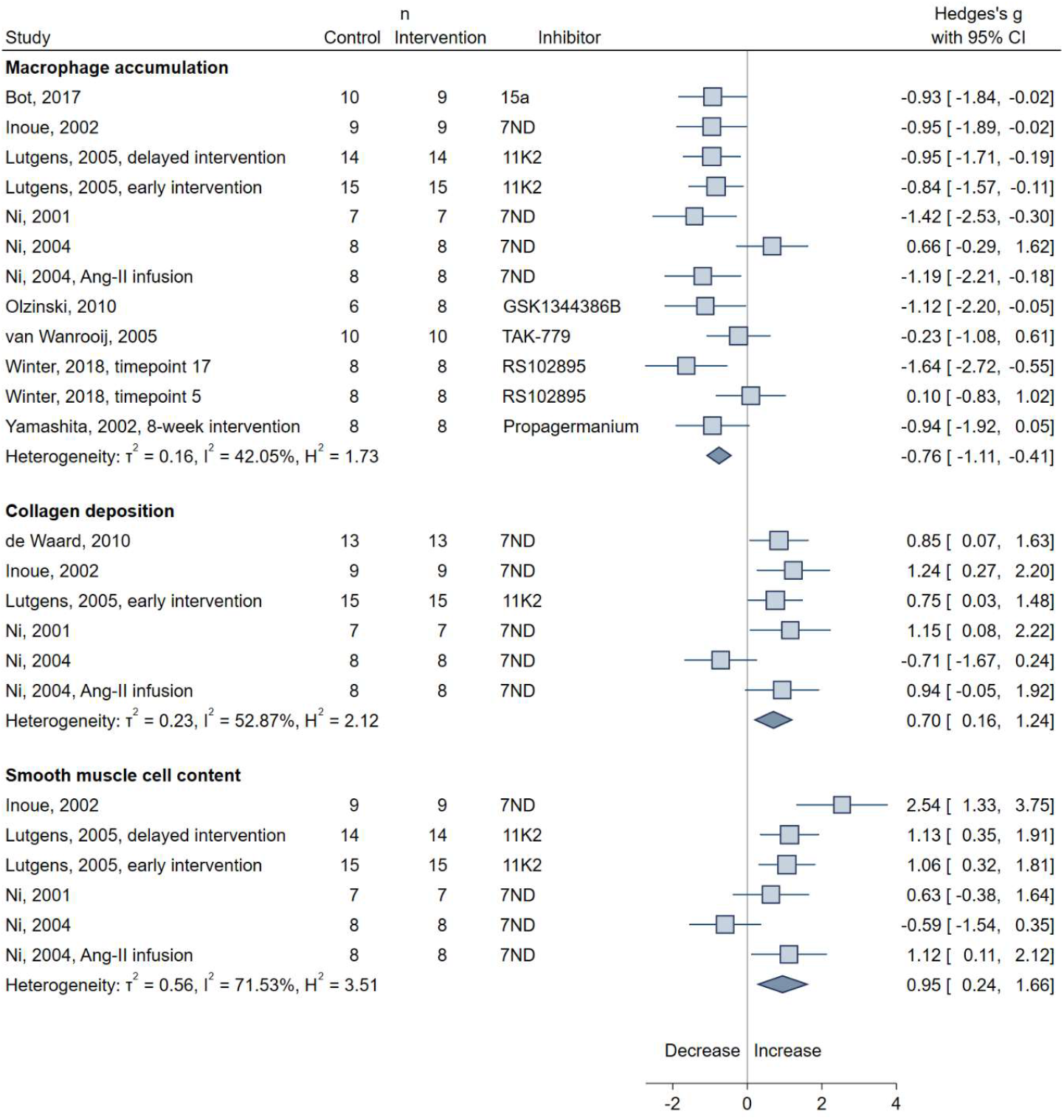
Forest plot of the effects of CCL2/CCR2 inhibition versus control (Hedges’ g) on macrophage accumulation, collagen deposition, and smooth muscle content in plaques of the aortic arch or root. Shown are the standardized mean differences, calculated as Hedges’ *g*, with their respective 95% confidence intervals per study. Plot squares are weighted for study size and correspond to individual effects, whereas plot whiskers correspond to the 95% confidence intervals. Diamonds indicate the pooled effects for each vascular bed. τ^2^, I^2^ and H^2^ as indicators of group heterogeneity are displayed for the groups containing more than one study.

Associations between CCL2/CCR2 inhibition and secondary outcomes are presented in **Online Figure I** and **Online Table I**. The experimental groups did not undergo changes in body weight, plasma triglycerides or blood monocytes. However, there was a significant increase in CCL2 plasma levels across studies inhibiting CCR2 and a significant decrease in IL-6 expression levels within plaques. There was a borderline association between CCL2/CCR2 inhibition and lower plasma total cholesterol levels.

### Subgroup analyses reveal no differences by lesion stage or intervention target

There was at least moderate heterogeneity (*I*^*2*^>50%) for all main outcomes except for macrophage accumulation (*I*^*2*^=42%). To explore whether other study variables could explain the between-study differences we performed a series of subgroup analyses (**Fig. 4** and **Online Table II**). There were no significant differences between subgroups of different stages of atherosclerosis progression (early, intermediate, advanced) at the time of onset of intervention, although there was a tendency for smaller effect sizes in mice with more advanced lesions. Similarly, we observed no difference in the effects of intervention on lesion size in the aortic arch or root between targets of intervention, with both CCL2 and CCR2 inhibition showing significant reductions. Lesion size reduction differed significantly between WTD-fed mice and mice fed chow (p=0.048) with the latter showing no significant reduction in lesion size. All but one study examining aortic lesions used *Apoe*^-/-^ models of atherosclerosis, but the single study using *Ldlr*^*-/-*^ mice also showed a significant reduction in lesion size. No significant differences in effects were detected between male- and female-specific analyses. None of the subgroup analyses resolved the heterogeneity between studies. A meta-regression analysis revealed an association between longer intervention duration and larger atheroprotective effects on lesion size (β=-0.153 [-0.285 to -0.021], p=0.023; **Online Figure IIA**), but failed to account for study heterogeneity (residual *I*^*2*^=67%).

**Figure 4.**
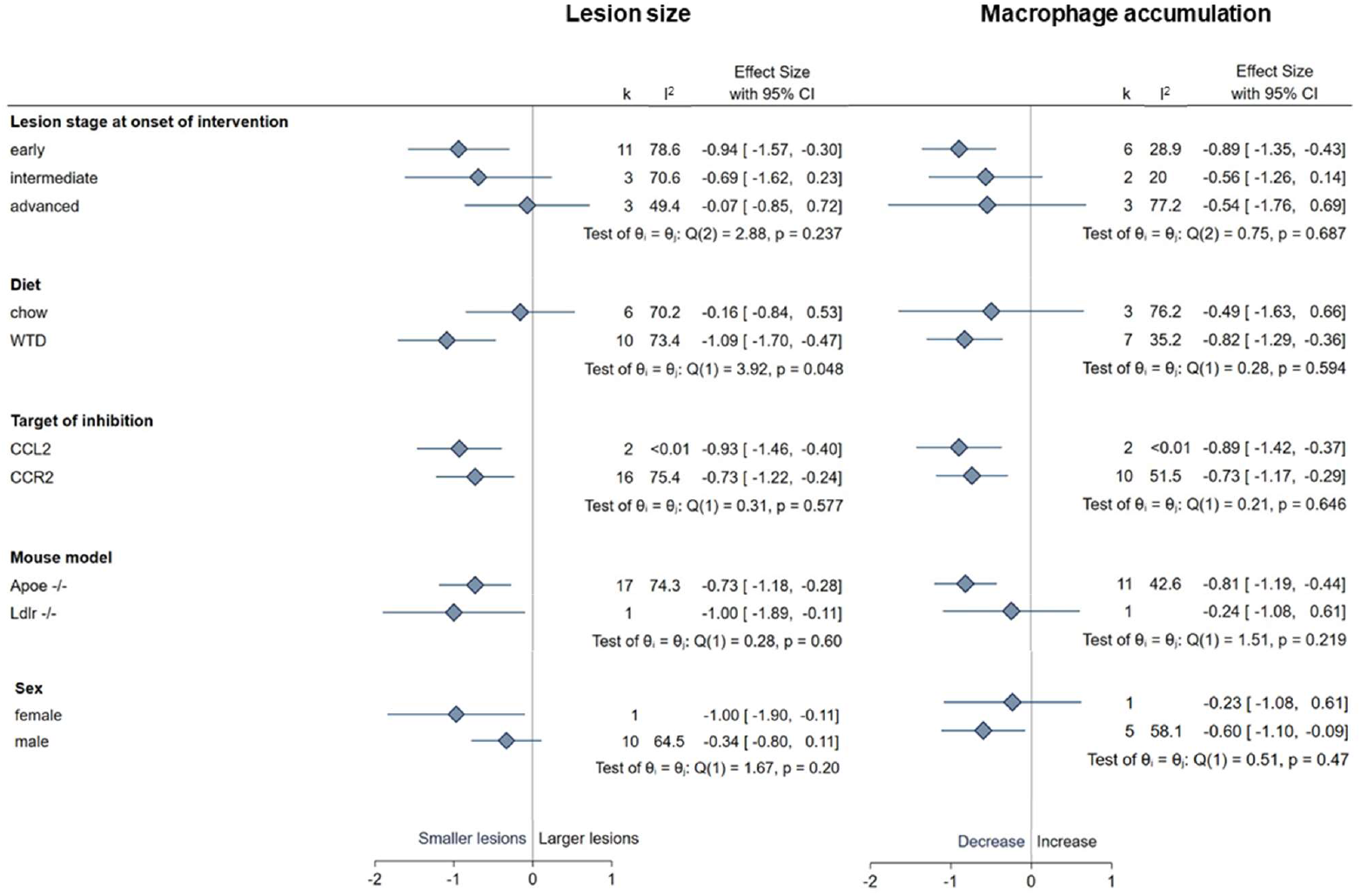
Subgroup analyses regarding the effects of CCL2/CCR2 inhibition versus control (Hedges’ g) on aortic plaque burden and macrophage accumulation by various study characteristics. Shown are the pooled standardized mean differences, calculated as Hedges’ *g*, with their respective 95% confidence intervals for each subgroup. Diamonds correspond to pooled effects per subgroup, whereas whiskers correspond to the 95% confidence intervals. Number of study arms (*k*) and heterogeneity measures (*I*^2^) per subgroup are displayed. The Cochran’s Q test and its p-value are provided as measures of between-subgroup differences.

### Effects of CCL2/CCR2 inhibition on intralesional macrophage accumulation predict the reductions in lesion size

To further explore sources of the derived heterogeneity, we performed meta-regression analyses exploring the associations between the effects of CCL2/CCR2 inhibition on features of plaque stability and their effect on lesion size. We found a significant association between the effects of different interventions on macrophage accumulation within plaques and the effects on the overall aortic lesion size (β=0.789 [0.263 to 1.314], p=0.003; **Fig. 5**). Residual heterogeneity (*I*^*2*^) after meta-regression was 27% compared to the initial 73%, thus invoking the notion that differences across the interventions in their effects on macrophage accumulation within plaques could explain 62% of the differences in overall effect on lesion size. There was no significant association between the effects of CCL2/CCR2 inhibition on plasma CCL2 levels and its effect on lesion size (**Online Figure IIB**).

**Figure 5.**
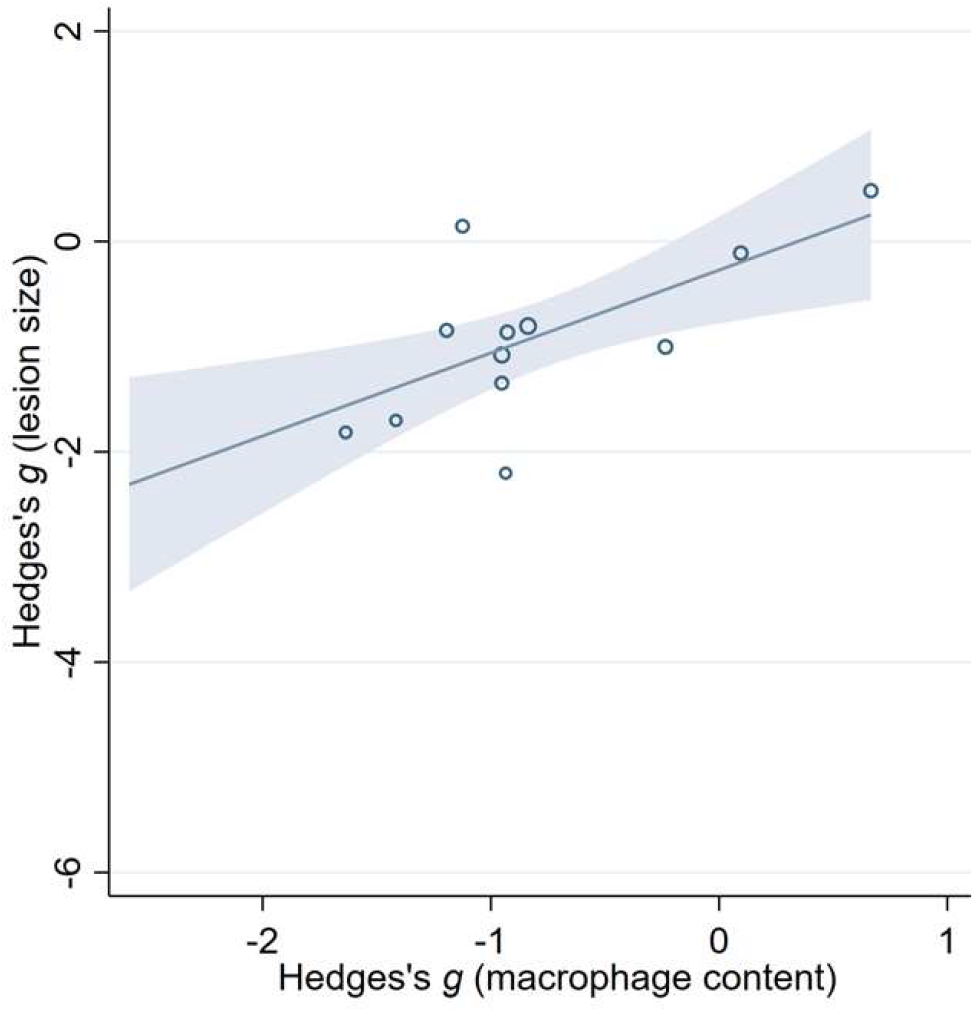
Meta-regression analysis of the effects of CCL2/CCR2 inhibition versus control (Hedges’ g) on macrophage accumulation on the effects of the intervention on atherosclerotic lesion size in the aortic arch and root. Data points indicate individual studies around the regression line with its 95% confidence interval (dotted lines).

### Publication bias and risk of bias assessment

Applying the Egger’s test,^60^ we detected a significant small-study effect (β=-7.95 [-12.08 to -3.82] p=0.0002) in the main analysis exploring the effects of CCL2/CCR2 inhibition on aortic lesion size, thus indicating presence of potential publication bias. After visual inspection of the respective funnel plot (**Online Figure III**), we explored whether a single outlier study^52^ could account for the observed effect. Following exclusion of this study, the observed small-study effect was attenuated (β=-5.79, [-11.77 to 0.19], p=0.058), while the overall effect of CCL2/CCR2 inhibition on aortic lesion size remained stable (g=-0.55, [0.93 to -0.17], p=0.005).

Finally, all eligible studies underwent a thorough quality assessment with the SYRCLE risk of bias tool.^34^ The detailed results are presented in **Fig. 6**. Importantly, there was evidence of high risk of detection bias in eleven eligible studies due to lack of blinding during outcome assessment. Furthermore, we detected high risk of attrition bias in eight studies, which failed to report exact sample sizes for every experiment or reasons for differing sample sizes across experiments of the same cohorts. All eligible studies were also assigned a high risk of reporting bias, because of the lack of a published pre-defined study protocol. The tool items referring to selection or performance bias, as well as the detection bias item for randomness of outcome assessment could not be assessed for most eligible studies due to insufficient information provided within the respective publications.

**Figure 6.**
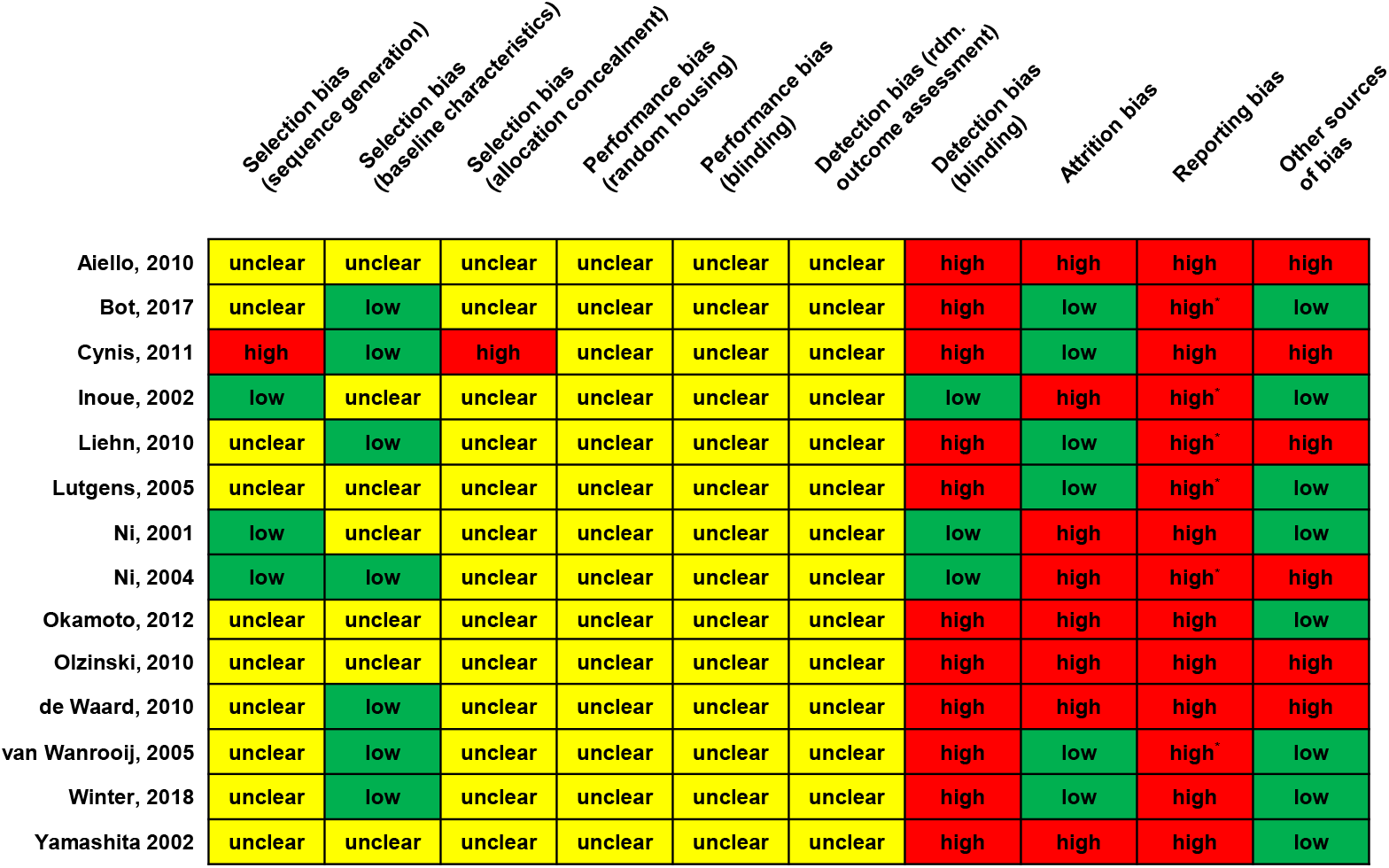
Assessment of risk of bias of included studies with the SYRCLE tool. * No study protocol was available, but key outcomes were all reported on. Key outcomes were pre-specified as lesion size plus two out of the following outcomes: Macrophage accumulation, collagen deposition, smooth muscle cell content.

## Discussion

Pooling data from 14 preclinical studies of experimental atherosclerosis in a comprehensive meta-analysis, we demonstrate that pharmacological blockade of CCL2 or CCR2 in mice leads to a significant reduction in atherosclerotic lesions in the aorta, the carotid, and the femoral artery. Furthermore, pharmacological inhibition of CCL2 or CCR2 causes reductions in intralesional macrophage accumulation and increases in plaque smooth muscle cell content and collagen deposition, consistent with a plaque-stabilizing effect. These effects are similar when targeting either CCL2 or CCR2, but were stronger for lesions in the carotid and femoral arteries than in the aorta. While there is substantial heterogeneity in the extent of CCL2/CCR2 inhibition on atherosclerotic lesion size, these effects were highly correlated with the effects of the interventions on macrophage accumulation within plaques, thus supporting the notion that intralesional macrophage reduction can serve as a surrogate marker of efficacy of CCL2/CCR2 inhibition.

The consistently large effects of pharmacological CCL2/CCR2 inhibition on atherosclerotic lesions across studies with different designs and across different vascular beds, when seen in conjunction with previous findings in *Ccl2*^*-/-*13^ or *Ccr2-/-*^12, 19^ mice testing the genetic deletion of the ligand or the receptor, provide strong preclinical support for the candidacy of CCL2/CCR2 signaling as a promising target in atherosclerosis. While data from clinical trials remain limited,^24,61^ recent results from genetic^20^ and epidemiological^21, 22^ studies emphasize a causal role of the CCL2/CCR2 axis in human atherosclerosis. Hence, there is coherent evidence from preclinical, genetic, epidemiological, and early-phase clinical trials that targeting the CCL2/CCR2 pathway may be a viable strategy to mitigate the risk of atherosclerotic disease.

Interestingly, we found stronger attenuating effects of CCL2/CCR2 inhibition on atherosclerotic lesions in the carotid artery, as compared to lesions in the aortic arch and root. While these findings cannot be directly translated to humans, they are consistent with the stronger associations we previously found between both genetically determined, as well as measured CCL2 circulating levels and risk of ischemic stroke, as compared to coronary artery disease and myocardial infarction.^20-22^ Despite the common mechanisms underlying atherogenesis and atheroprogression across different vascular territories, differences in the effects of established risk factors, such as smoking and hypertension, on atherosclerotic manifestations from different vascular beds are well-known.^62-64^ Whether pharmacological targeting of the CCL2/CCR2 axis and inflammation in humans differentially affects atherosclerotic lesion formation in different vessels would need to be further explored in future studies. Still, this could have implications for the selection of the right population for future clinical trials.

Aside from its influence on lesions size, CCL2/CCR2 inhibition further exerted a stabilizing effect on plaques. Specifically, mice in the intervention arms exhibited a lower macrophage accumulation, and a higher smooth muscle cell and collagen content, consistent with a smaller core and a thicker fibrous cap, both characteristics of a plaque less vulnerable to rupture and subsequent complications like acute coronary syndromes or ischemic stroke.^65, 66^ Our results are consistent with those of a recent cross-sectional study of plaque samples from patients undergoing carotid endarterectomy, which showed associations between CCL2 levels within plaques and histopathological features of plaque vulnerability.^23^ Thus, these data support a role of CCL2/CCR2 beyond the early stages of atherogenesis and highlight the potential benefits of targeting this axis even in patients with established atherosclerotic disease in future trials.

In a meta-regression analysis, we found the heterogeneity of the effects of CCL2/CCR2 inhibition on aortic lesion size to be to a large extent explained by the effects on plaque macrophage accumulation. This observation agrees with the key role of the CCL2/CCR2 axis in attracting monocytes to the atherosclerotic lesion,^67^ but also indicates that the effects of pharmacological approaches targeting the CCL2/CCR2 axis on intralesional macrophage accumulation could be used as a surrogate marker of the overall efficacy of CCL2/CCR2 inhibition. The latter could have implications for the design of future early-phase clinical trials and the identification of a proper readout for drug response and efficacy beyond clinical endpoints that would require very large sample sizes.

Another important finding of the current meta-analysis is the lack of heterogeneity in efficacy between studies targeting either CCL2 or CCR2. The consistency in the effects of molecules targeting either the ligand CCL2 or its receptor CCR2 for either decreasing lesion size or improving the plaque stability profile indicate no superiority of one approach over the other in animal models of atherosclerosis. This is important given the different structural and targeting properties of the pharmaceutical agents employed. Features such as surface coverage, binding affinity, or bioavailability differ substantially between antibodies, orthosteric small molecule inhibitors, or decoy ligands. Moreover, most of the agents included in our meta-analysis were developed before the CCR2 X-ray structure was available.^68^ Also, it should be noted that there are fewer agents targeting chemokine ligands, as compared to chemokine receptors,^9, 69^ which is also reflected in the low number of studies inhibiting CCL2 in our meta-analysis. Moving towards clinical trials in humans, it would be important to consider both approaches as potentially promising.

Our study has specific methodological strengths. Incorporating data from studies applying different approaches allowed us to explore at a meta-analysis level the comparativeness of different approaches targeting either CCL2 or CCR2 and offered us sufficient power to detect a clear plaque-stabilizing effect of CCL2/CCR2 inhibition beyond a reduction in lesion burden, as well as differential effects across different vascular territories. Furthermore, using meta-regression analyses, we were able to identify a correlation between the effects of CCL2/CCR2 inhibition on plaque macrophage accumulation and plaque lesion size, thus also offering mechanistic insights into the atheroprotective effects of CCL2/CCR2 inhibition.

Our study also has limitations. First, there was considerable between-study heterogeneity in almost all analyzed outcomes, which could bias the derived effect estimates. It is possible that differences in experimental design as well as in the efficacy of the tested interventions underly this heterogeneity. Still, heterogeneity could not be resolved in any of the subgroup analyses. Second, there was evidence of small-study effects indicating publication bias. While publication bias could indeed influence the effect estimates, we found the small-study effect to be primarily driven by a single outlier study. Reassuringly, the effects were only slightly attenuated after exclusion of this study from the meta-analysis. Third, in our risk of bias assessment, we found the majority of the included studies to fulfill few of the quality criteria and to be vulnerable to detection, attrition, and reporting bias. This necessitates cautious interpretation of the findings, as sources of bias in preclinical studies can contribute to lack of translation of promising preclinical experiments into successful clinical trials.^70-72^ Fourth, some of our analyses, such as the analyses for lesions in the carotid and femoral arteries, the analyses for CCL2 inhibition, the subgroup analyses per stage of atherosclerotic lesions, and some meta-regression analyses were based on a rather small number of study arms and are thus limited by low statistical power. Fifth, the lesion staging used in the subgroup analysis relied on the age of mice, feeding, and treatment durations rather than histopathological lesion assessment due to paucity of data. Lastly, agents used in some studies, like 7ND, appear impractical for therapeutic approaches in humans compared to small molecule inhibitors.

In conclusion, our findings demonstrate a clear atheroprotective effect of the pharmacological targeting of the CCL2/CCR2 axis in mouse models of atherosclerosis. Our meta-analysis supports a comparable efficacy of approaches targeting either CCL2 or CCR2, differences in efficacy across vascular territories, stabilizing effects on plaque morphology beyond reductions in lesion burden, and a mediating role of intralesional macrophage accumulation in the atheroprotective effects of CCL2/CCR2 inhibition. This preclinical evidence, when seen along with recent data from human studies, highlights the translational potential of targeting CCL2/CCR2 signaling in atherosclerotic disease and provides important insights for informing the design of future clinical trials.

### Sources of funding

This study has received support by the Deutsche Forschungsgemeinschaft (DFG) through grants for the Collaborative Research Center 1123 (CRC1123, L.Z., fellowship IRTG1123; Y.A., project B03; J.B. project A03; M.D. project B03). L.Z. received a graduate scholarship from the Medical Faculty of the Ludwig-Maximilians-University (LMU) Munich. Y.A. was supported by grants from the FöFoLe program of LMU Munich (FöFoLe 921) and the Friedrich Baur-Stiftung. M.G. received funding from the Onassis Foundation and the German Academic Exchange Service (DAAD).

### Disclosures

None.

## Non-standard Abbreviations and Acronyms

CCL2: C-C chemokine ligand 2
CCR2: C-C chemokine receptor 2
CCR5: C-C chemokine receptor 5
CXCR3: C-X-C chemokine receptor 3
IL-1β: Interleukin 1 beta
IL-6: Interleukin 6
TNF-α: Tumor necrosis factor alpha

## Notes

### Competing Interest Statement

The authors have declared no competing interest.

