## Supplemental Material for "CCL2/CCR2 inhibition in atherosclerosis: a meta-analysis of preclinical studies"

### ONLINE SUPPLEMENTAL MATERIAL

**Online Table I. Summary effects of CCL2/CCR2 inhibition on secondary outcomes.** Shown are overall Hedges' g, along with 95% confidence interval and p-value, as derived from random-effect meta-analyses. Heterogeneity is expressed as $I^2$, and the number of study arms included in meta-analysis with *k*. * Significant at $p < 0.05$.

| | **Body weight** | **Total plasma cholesterol** | **Total triglycerides** | **Blood monocytes** |
|---|---|---|---|---|
| *g* | 0.17 | -0.37 | 0.15 | 0.04 |
| [95% CI], p | [-0.34; 0.67], 0.522 | [-0.66; -0.08], 0.013* | [-0.17; 0.48], 0.363 | [-0.86; 0.93]; 0.938 |
| $I^2$ | 47.4% | <0.01% | <0.01% | 75.5% |
| *k* | 6 | 10 | 8 | 5 |

| | **Plasma CCL2** | **CCR2 mRNA** | **IL-6 mRNA** | **TNF-α mRNA** |
|---|---|---|---|---|
| *g* | 1.29 | 0.71 | -1.06 | -0.67 |
| [95%-CI], p | [0.98; 1.61], $< 1\times10^{-5}$* | [-0.90; 2.31], 0.384 | [-2.04; -0.08], 0.035* | [-1.39; 0.05], 0.069 |
| $I^2$ | <0.01% | 84.9% | 50.8% | 18.4% |
| *k* | 9 | 3 | 3 | 3 |

**Online Table II. Subgroup analyses regarding the effects of CCL2/CCR2 inhibition versus control (Hedges' g) on collagen and smooth muscle cell (SMC) content in aortic lesions.** Shown are the pooled standardized mean differences, calculated as Hedges' *g*, with their respective 95% confidence intervals for each subgroup. Number of study arms (*k*) and heterogeneity measures ($I^2$) per subgroup are displayed. The Cochran's Q test and its p-value are provided as measures of between-subgroup differences. * Significant at p<0.05.

| | | **Collagen content** | | | **SMC content** | | |
|---|---|---|---|---|---|---|---|
| | **Subgroups** | *g* | [95% CI], p | $I^2$ (%)<br>*k* | *g* | [95% CI], p | $I^2$ (%)<br>*k* |
| **Lesion stage at intervention** | early | 0.89 | [0.28; 1.48], 0.003* | <0.01<br>2 | 0.91 | [0.31; 1.51], 0.003* | <0.01<br>2 |
| | intermediate | 1.00 | [0.40; 1.60], 0.001* | <0.01<br>2 | 2.54 | [1.33; 3.75], 0.0004* | (-)<br>1 |
| | delayed | 0.11 | [-1.51; 1.73], 0.897 | 82.0<br>2 | 0.25 | [-1.43; 1.93], 0.764 | 83.0<br>2 |
| | $Q_b$; *p* | | 1.03; 0.597 | | | 6.79; 0.030* | |
| **Diet** | chow | 0.48 | [-0.71; 1.68], 0.430 | 78.1<br>3 | 0.99 | [-0.76; 2.74], 0.267 | 88.0<br>3 |
| | WTD | 0.96 | [0.33; 1.59], 0.003* | <0.01<br>2 | 0.63 | [-0.38; 1.64], 0.219 | (-)<br>1 |
| | $Q_b$; *p* | | 0.47; 0.494 | | | 0.12; 0.728 | |
| **Target** | CCL2 | 0.75 | [0.03; 1.48], 0.041* | (-)<br>1 | 1.10 | [0.56; 1.64], <0.0001* | <0.01<br>2 |
| | CCR2 | 0.69 | [0; 1.38], 0.050 | (-)<br>5 | 0.89 | [-0.34; 2.11], 0.155 | 82.03<br>4 |
| | $Q_b$; *p* | | 0.02; 0.897 | | | 0.09; 0.76 | |
| **Model** | *Apoe*$^{-/-}$ | 0.68 | [0.14; 1.21], 0.013* | 52.8<br>6 | 0.95 | [0.24; 1.66], 0.009 | 71.5<br>6 |
| | *Ldlr*$^{-/-}$ | | (-) | | | (-) | |
| | $Q_b$; *p* | | (-) | | | (-) | |

**Online Figure I. Forest plots of the effects of CCL2/CCR2 inhibition versus control (Hedges' *g*) on secondary outcomes.** Shown are the results for **(A)** mouse body weight, **(B)** plasma cholesterol and **(C)** triglycerides, **(D)** circulating blood monocytes, **(E)** circulating CCL2, as well as tissue mRNA expression of **(F)** CCR2, **(G)** IL-6, and **(H)** TNF-α. Individual study arms' Hedges' *g* is provided alongside 95% confidence interval (95% CI) in brackets. Weighted plot squares indicate Hedges' *g*, whiskers depict the confidence interval.

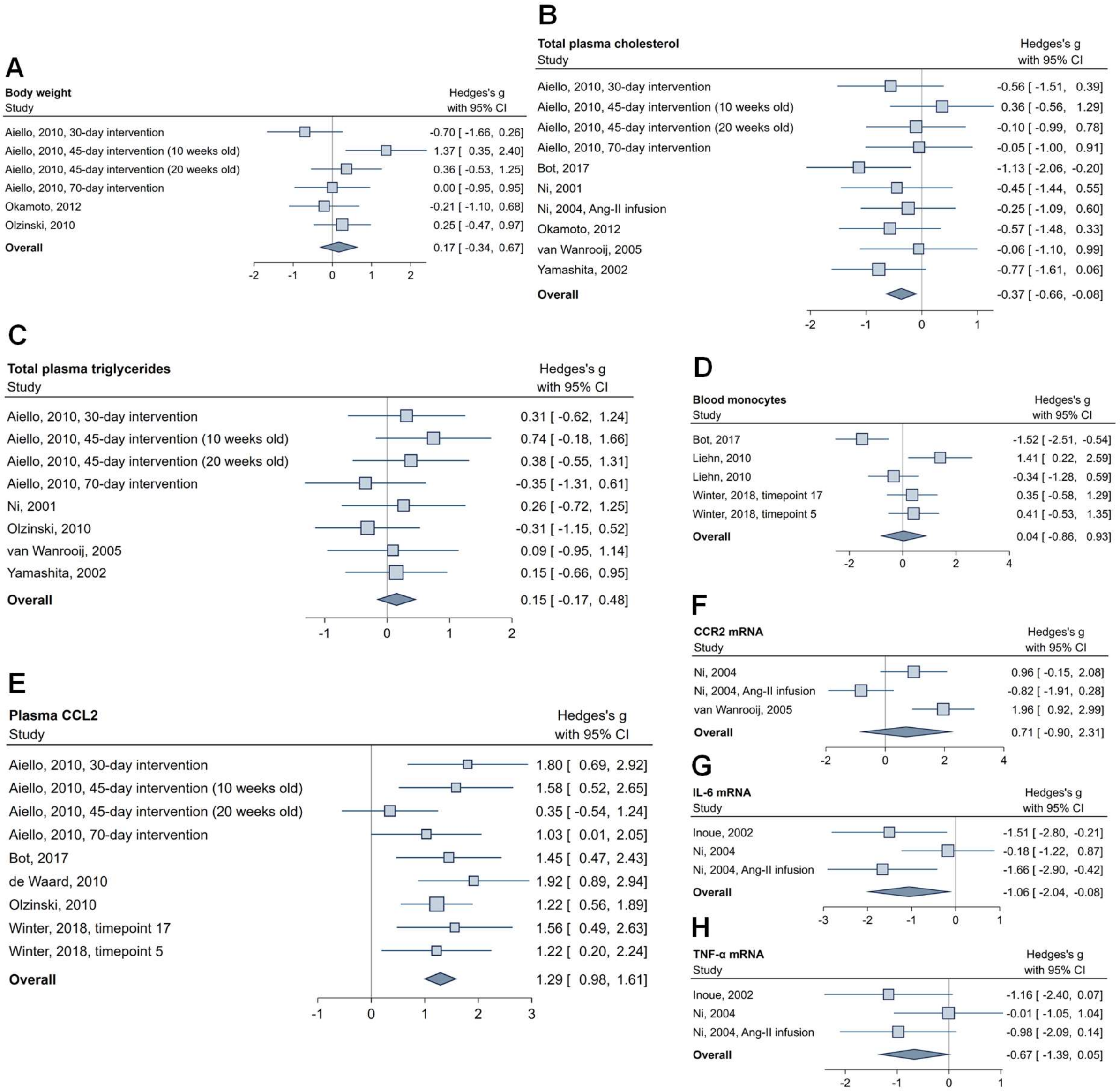

**Online Figure II. Meta-regression analysis of (A) duration of intervention and (B) the effects of the intervention on plasma CCL2 levels on the effects of the intervention on atherosclerotic lesion size in the aortic arch and root.** Predicted regression lines with 95% confidence interval are depicted for lesion size in the aortic arch/root against duration of intervention in weeks (**A**; (β=-0.153, 95%CI=[-0.285; -0.021], p=0.023, residual $I^2$=67.4%), and plasma CCL2 levels at end of intervention (**B**; β=0.634, 95%CI=[-1.248; 2.517], p=0.509, residual $I^2$=65.1%). Data points indicate individual studies around the regression line with its 95% confidence interval (blue shaded area).

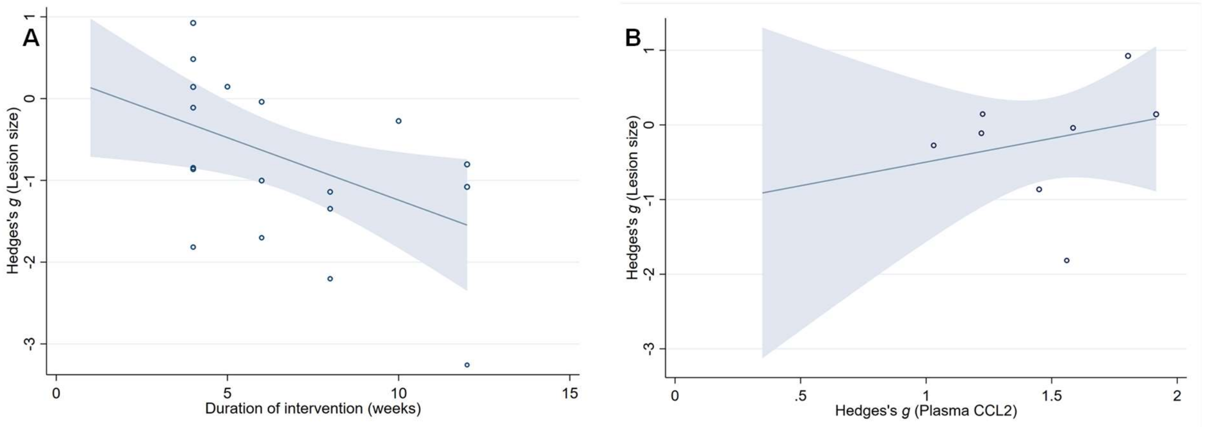

**Online Figure III. Funnel plot of the effects of CCL2/CCR2 inhibition versus control (Hedges' g) in the individual studies on atherosclerotic lesion size in the aortic arch or root.** Circles indicate individual studies. A standard-error-adjusted pseudo-95% confidence range is indicated with dotted lines. The vertical blue line corresponds to the pooled Hedges' *g* for lesion size in the random-effects meta-analysis.

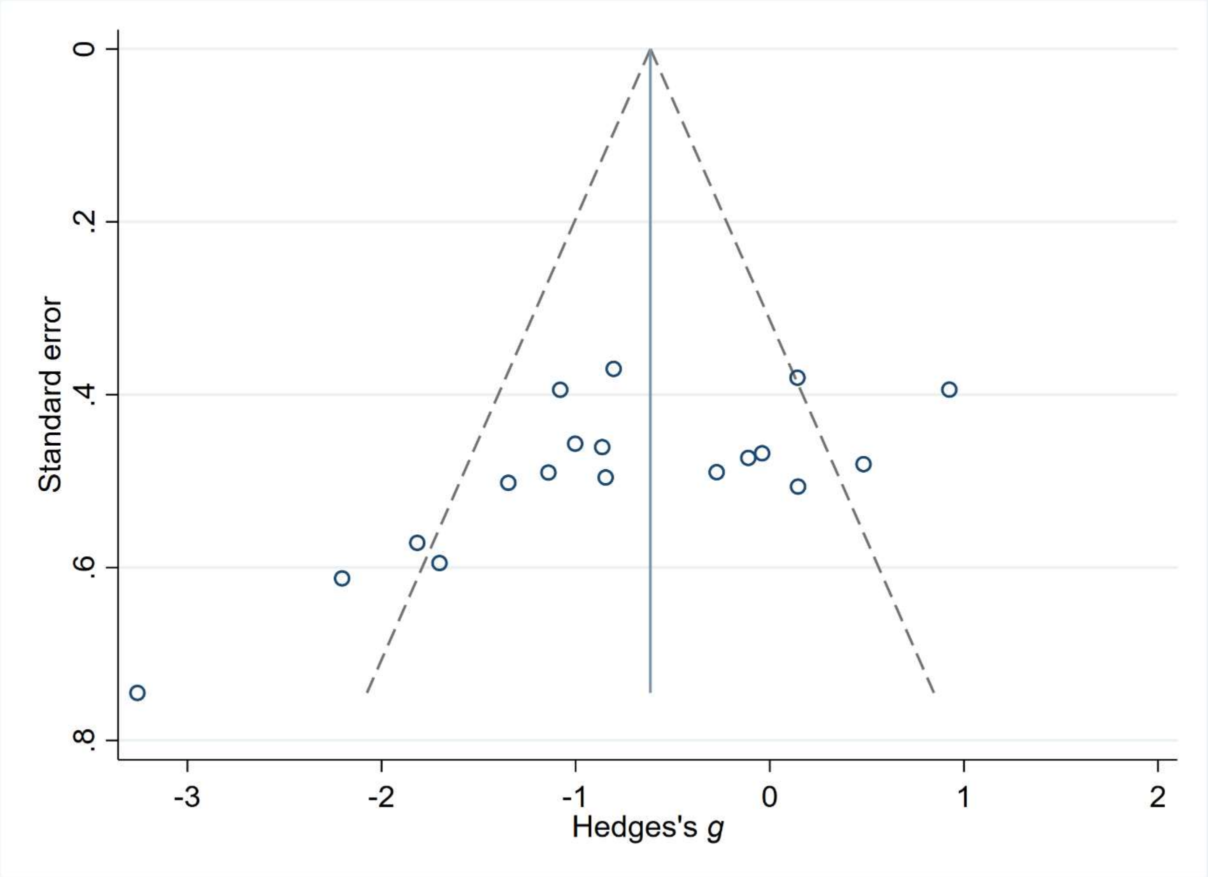
